# EOR-1/PLZF-regulated WAH-1/AIF sequentially promotes early and late stages of non-apoptotic corpse removal

**DOI:** 10.1101/2024.12.04.626465

**Authors:** Nathan Rather, Melvin Williams, Aladin Elkhalil, Karen Juanez, Rashna Sharmin, Ginger Clark, Shai Shaham, Piya Ghose

## Abstract

Programmed cell death (PCD) is a crucial, genetically-encoded, and evolutionarily-conserved process required for development and homeostasis. We previously identified a genetically non-apoptotic, highly ordered, and stereotyped killing program called Compartmentalized Cell Elimination (CCE) in the *C. elegans* tail-spike epithelial cell (TSC). Here we identify the transcription factor EOR-1/PLZF as an important coordinator of CCE. Loss of EOR-1 results in a large, persisting, un-engulfed soma with enlarged nuclei. We find that EOR-1 and its partners positively regulate the transcription of the Apoptosis Inducing Factor AIF homolog, WAH-1/AIF. We report stereotyped and sequential spatiotemporal dynamics of WAH-1/AIF1 during phagocytosis, with defined roles acting early and late, within the dying cells. Mitochondria to plasma membrane translocation within the TSC soma is required its internalization by its phagocyte, and plasma membrane to nuclear translocation is required for DNA degradation and ultimately, corpse resolution. Our study suggests that EOR-1 serves as a master regulator for the transcriptional control of DNA degradation is essential for changes in nuclear morphology required for cellular dismantling and infers that tight spatiotemporal regulation of WAH-1/AIF is required for this function.

**Summary Statement:** This work describes the genetic control and cellular dynamics of a factor linked to cancer, metabolic and degenerative disease acting in developmentally dying cells to instruct their own removal.

## Introduction

Programmed cell death (PCD) is a genetically-encoded and evolutionarily-conserved cell elimination vital for normal development and homeostasis (1), with defects linked to disease (2–4). Apoptosis, the most well-studied form of PCD, is by defined by cell shrinkage, chromatin condensation, nuclear fragmentation, and mitochondrial fragmentation (1, 5, 6). Genetically, apoptosis is controlled by several conserved regulators, including the cysteine protease caspase family. Caspases have a conserved pathway of regulators, including the pro-apoptotic Apaf-1, the antiapoptotic Bcl-2, and the proapoptotic BH3-only in mammals (7–9). Several forms of non-apoptotic cell death have been also described (10).

In addition to the cell killing, cell remains must be efficiently removed via phagocytosis to avoid secondary necrosis and autoimmune consequences (11, 12). The mechanism of apoptotic corpse phagocytosis is well described. First, the dying cells are recognized (13), and engulfed (14); once fully internalized and encapsulated in a phagocytic vesicle called a phagosome, the corpse is eventually resolved through multiple steps of increased acidification of the phagosome (15) and introduction of lysosomal hydrolases (16). Apoptotic engulfment and phagosome maturation pathways are conserved in mammals (17, 18).

Much of what we know about apoptotic cell killing and clearance has been gleaned using the powerful genetic model organism, the nematode *C. elegans* (19) (20). Phagocytosis has also been extensively characterized in the nematode, with seminal work conducted on the molecular machinery and cell biology required for corpse engulfment (21) and phagosome maturation (22). Newer studies have described the role of engulfment machinery genes in neuronal sculpting and function (23), in polar body removal (24), and in cell clearance following LCD and other deaths (25, 26).

We previously described the molecular pathway required for the Compartmentalized Cell Elimination (CCE) of the tail-spike epithelial scaffolding cell (TSC) in the *C. elegans* embryo, a tripartite cell killing program required for the death and clearance of complex cells (27). Briefly, CCE entails three distinct and stereotyped cell elimination events. As the CCE program occurs 10 times slower than most apoptotic nematode cells, it allows for detailed examination of these events at the gross and subcellular level (27). During CCE, the single process of the TSC segments into two morphologically distinct sub-compartments that exhibit degenerative morphologies resembling two different forms of neurite pruning (27). The binucleate soma rounds, superficially resembling apoptotic morphology (27). The highly specialized morphology of the TSC allows it to serve as a timekeeper for the steps of CCE.

Molecularly, CCE is genetically distinct from apoptosis (28–30). While it does require the *C. elegans* caspase CED-3, it is not regulated by the BH3-only homolog EGL-1 (30), and additional non-canonical regulators are required for cell killing (28–30) During engulfment, the canonical engulfment machinery operates only on the soma corpse (27). Dismantling of the TSC process remnants instead require the cell-cell fusogen EFF-1 for phagosome sealing (27).

In the present study, we identify EOR-1/PLZF as the first transcriptional regulator of a cell clearance regulator, the homolog of the Apoptosis inducing factor WAH-1. We characterize WAH-1 dynamics and subcellular localization and define dual roles in sequential steps of cell corpse removal. We find that EOR- 1-regulated WAH-1 acts in the dying TSC to promote the careful timing of both corpse internalization and resolution. Failure of EOR-1/PLZF and/or WAH-1/AIF function leads to a corpse progression transition state, characterized by lack of cellular engulfment and failure of lysosomal corpse degradation.

## Results

### The transcription factor EOR-1/PLZF promotes CCE compartment-specifically

The TSC **(Figure 1A**) extends a single posteriorly-directed process when intact (IMA, intact, mature). During early CCE (BA, beading, attached; BD, beading, detached), this single process shows axon pruning-like morphology proximally and axon retraction-like pruning distally (**Figure 1B**). At this step, the binucleate soma rounds in a manner superficially resembling apoptotic cell death. Later in CCE, the distal process retracts into a distal node (**Figure 1C**) (soma, distal retraction, SDR) leaving behind a soma and a distal remnant (soma, distal degrading, SDD) (**Figure 1D**) which are phagocytosed stochastically by different neighboring phagocytes before the animal hatches. The highly organized and stereotyped elimination sequence of CCE allows us to use the cell’s general morphology as a timekeeper for events in the cell death program.

**Figure 1.**
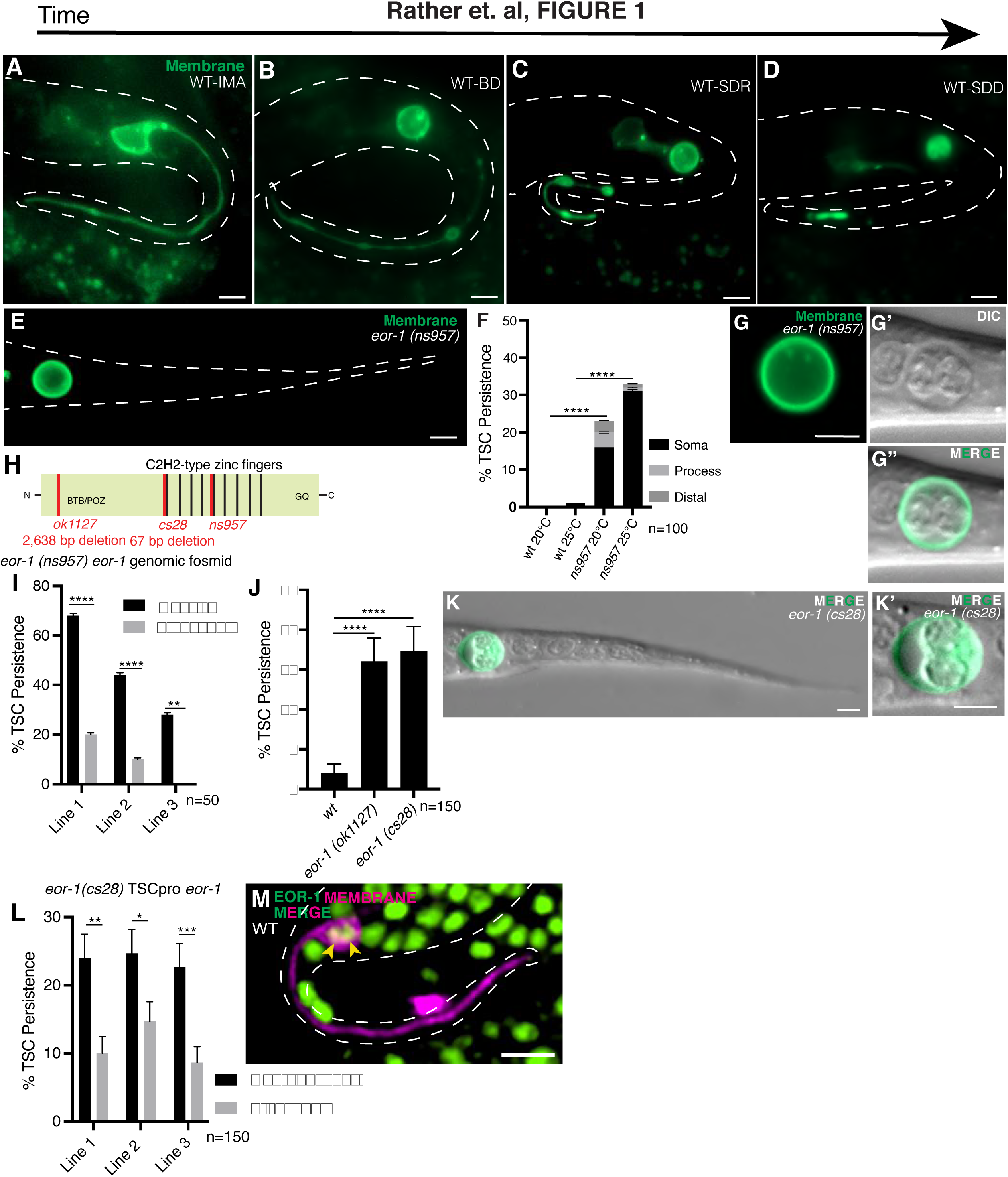
EOR-1/PLZF promotes CCE cell-autonomously. (A-D) Stages of CCE. IMA, intact mature cell; BD, beading, detached; SDR, soma-distal retracted; SDD, soma, distal degrading. n = 10 biologically independent animals each with similar results. **(E)** Mutant *ns957* at L1 stage showing persisting TSC soma. n = 10 biologically independent animals each with similar results **(F)** Quantification of *ns957* CCE defects. Data are mean +/_ s.e.m. Statistics: two-tailed unpaired student’s t-test, see Supplementary Table 3 for individual P values. **(G-G’’)** Enlarged view of *ns957* persisting soma to show DIC view. **(I)** Gene structure of *eor-1*. **(J)** Quantification of *eor-1* fosmid DNA rescue of *ns95*7, n=50. (**K)** Quantification CCE defects of other *eor-1* alleles, n=150. **(L, L’)** Quantification of CCE defects of additional *eor-1* mutant alleles. **(M)** *eor-1* TSC-specific rescue, n=150. **(N)** *eor-1* expression, *eor-1* promoter-driven mKate2. n = 10 biologically independent animals each with similar results. Scale bar, 5 μm. Data are mean +/ s.e.m. Statistics: two-tailed unpaired student’s t-test, see Supplementary Table 3 for individual P values.

While CCE is dependent on the main *C. elegans* caspase CED-3 (27) it is genetically distinct from apoptosis as it does not require EGL-1/BH3-only (30) CCE is also temporally distinct from apoptosis as it takes longer to complete. While most cells fated to die apoptotically in *C. elegans* live for only 30 minutes (31), the TSC lives for 5 hours (32) with CCE of the TSC taking 150 minutes (27). This longer timescale positions CCE of the TSC as a promising platform to better understand the fundamental cell biology of programmed cell elimination in greater detail.

We performed a forward genetic screen for CCE defects of the TSC using a membrane marker strain for the TSC (TSC-myrGFP) that we have previously employed (27). We obtained a mutant, *ns957,* bearing an inappropriately persisting and enlarged cell body only, but with no process **(Figure 1E)**. This CCE defect increased when animals were raised and scored at 25°C when compared to 20°C **(Figure 1F)**.

Intrigued by a mutant where only soma elimination fails but where the process is successfully eliminated, we examined the morphology of our mutant more carefully using Differential Interference Contrast (DIC) microscopy. Apoptotic cell death has well-documented morphological characteristics that are visible under DIC microscopy, such as pyknotic nuclei and refractility (33). However, we observed in our *ns957* mutant enlarged and non-pyknotic nuclei, and a non-refractile cytoplasm (**Figure 1G-G’’**).

Following Whole Genome Sequencing, genomic DNA rescue (**Figure 1I**), and examination of other alleles (**Figure 1J, K-K’**), we found that our screen mutant has a causative lesion in the gene encoding the transcription factor EOR-1/PLZF, resulting in an E570K change in the Zn finger domain (**Figure 1H**). We also found from cell-specific rescue experiments that *eor-1* acts cell-autonomously in the TSC (**Figure 1L**) and that *eor-1* is expressed in the nucleus, consistent with the notion that it acts as a transcription factor (**Figure 1M**).

EOR-1 is the ortholog of the BTB/zinc-finger tumor-suppressor transcription factor PLZF and has been reported to be a regulator for both apoptotic and non-apoptotic (LCD) forms of PCD identified in *C. elegans* (34). In a non-cell death context, EOR-1 is involved in neuronal specification (35). However, in terms of PCD, EOR-1/PLZF has been implicated both in the apoptotic deaths of the HSN neurons (34) as well as in non-apoptotic Linker-type Cell Death (LCD) (36). EOR-1 appears to play a specification role upstream to the execution phase of these deaths, acting upstream or in parallel to CED-9 in the HSNs and upstream of HSF-1 in conjunction with WNT signaling in LCD (34). However, its precise mechanistic contribution to either form of PCD remains unknown.

### EOR-1/PLZF acts downstream of CED-3/Caspase, but with known regulatory partners

We proceeded to examine the precise role of EOR-1 in PCD using CCE of the TSC as our model. In its known roles in cell fate, EOR-1 functions with known co-regulators of transcription. We first examined a role of EOR-2, which binds to the BTB domain of EOR-1 and promotes its function, (37) by looking at *eor-2 (cs42)* mutants (**Figure 2A, B, C**) and found that this phenocopied *eor-1* mutants. EOR- 1/PLZF has also been reported to act as a pioneer transcription factor that functions in conjunction with the MAU-2 cohesin loader, and the SWSN-1 SWI/SWNF chromatin remodeling complex member to modify chromatin architecture and gene expression of genes required for the maturation of the post-mitotic development of HSN neurons (35). To determine whether EOR-1 functions with MAU-2 and SWSN-1 to promote CCE, we examined *mau-2 (qm160)* **(Figure 2A, D, E)** and *swsn-1 (ku355)* **(Figure 2A, F, G)** mutant larvae; we found that they show persisting TSC somas that phenocopy the distinct morphology of *eor-1* mutant animals. Therefore, EOR-1 functions in the new setting of CCE via its previously identified partners in transcription.

**Figure 2.**
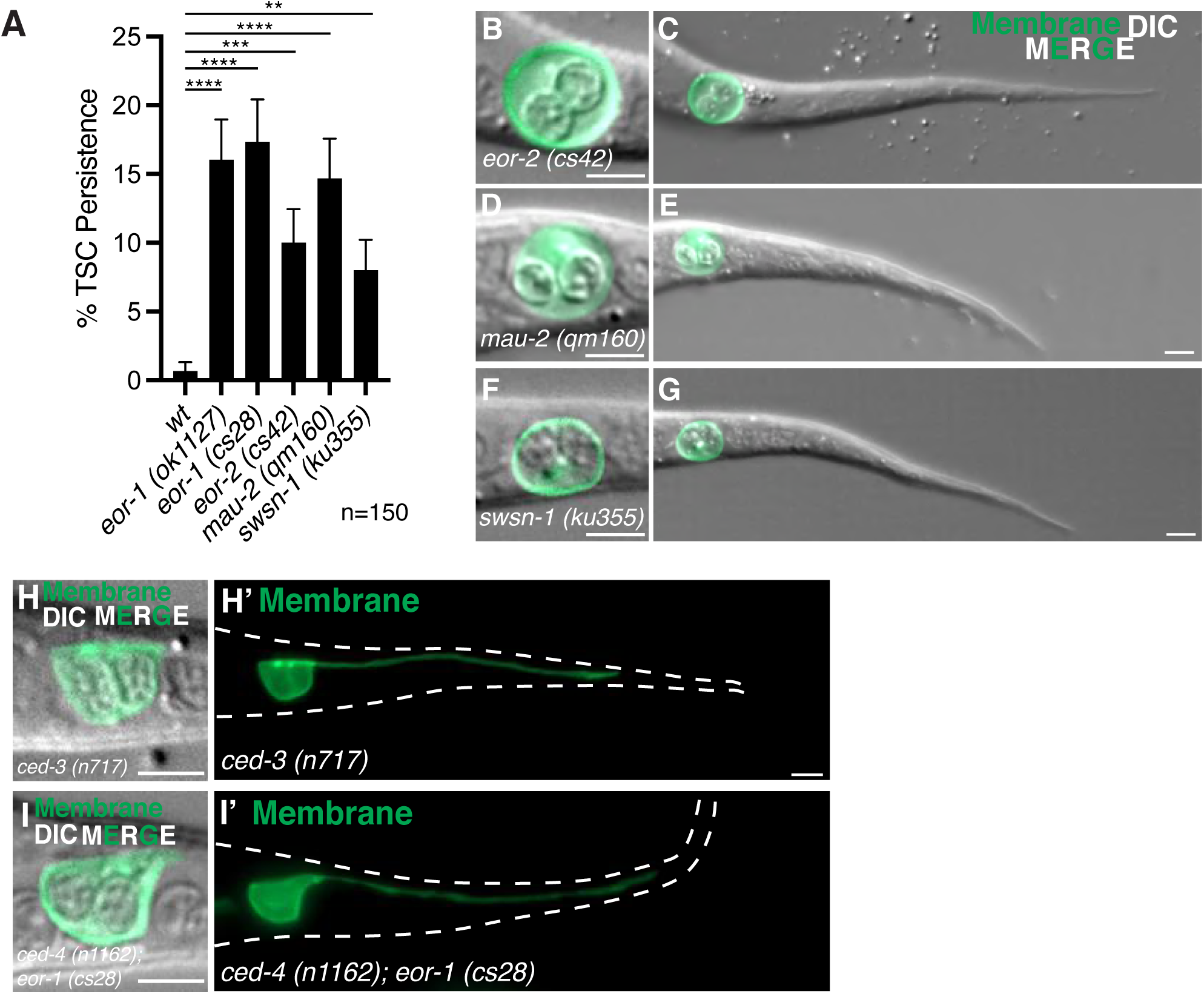
EOR-1/PLZF functions in CCE with its known partners but likely downstream of CED- 3/Caspase. **(A)** CCE defects across *eor-1* alleles and *eor-2*, *mau-2* and *swsn-1* mutants. Persisting TSC soma phenotypes in **(B, C)** *eor-2(cs42)*, **(D, E)** *mau-2 (qm160)* n = 10 biologically independent animals each with similar results and **(F, G)** *swsn-1 (ku355)* n = 5 biologically independent animals each with similar results. The TSC remains intact at L1 in both *ced-3 (n717)* **(H)** and *ced-4 (n1162); eor-1(cs28)* **(I)** n = 10 biologically independent animals each with similar results. Scale bar, 5 μm. n=150 for statistics. Data are mean +/_ s.e.m Statistics: two-tailed unpaired student’s t-test, see Supplementary Table 3 for individual P values.

We next asked whether EOR-1 functions upstream or downstream of CED-3/caspase and whether it impacts TSC fate in a more general way. As in the case of apoptosis, CCE requires the main *C. elegans* caspase CED-3, as well as its upstream regulator CED-4/Apaf1(30). We generated a *ced- 4(n1162); eor-1(cs28)* double mutant and found an intact TSC phenotype and nuclear phenotype characteristic of *ced-3(n717)* single mutants (N=10/10), suggesting that EOR-1 acts downstream of CED- 4 and CED-3 to promote CCE and likely does not affect TSC development **(Figure 2H-H’, I-I’).**

### EOR-1/PLZF is important late in CCE for corpse internalization

Given the striking soma morphology of *eor-1* mutants, we examined closely the soma morphology changes across the different CCE stages in both wild-type and *eor-1 (cs28)* mutants using a combination of DIC optics and our fluorescent TSC membrane reporter, and noted TSC soma features at different CCE stages. Prior to CCE, when the cell has matured, but is still intact (Intact mature stage, IMA) (**Figure 3A-B’’**), the soma appears polygonal with its nuclei appearing ovoid, flat and speckled with many small refractile bodies (**Figure 3B’-B’’**). As CCE sets in, the soma becomes round and the proximal process displays beading (Beading attached stage, BA) (**Figure 3C-D’’**), during which the nuclei appear round and spheroid (**Figure 3D’-D’’**). Nuclear contents retain a speckled appearance of refractile material.

**Figure 3.**
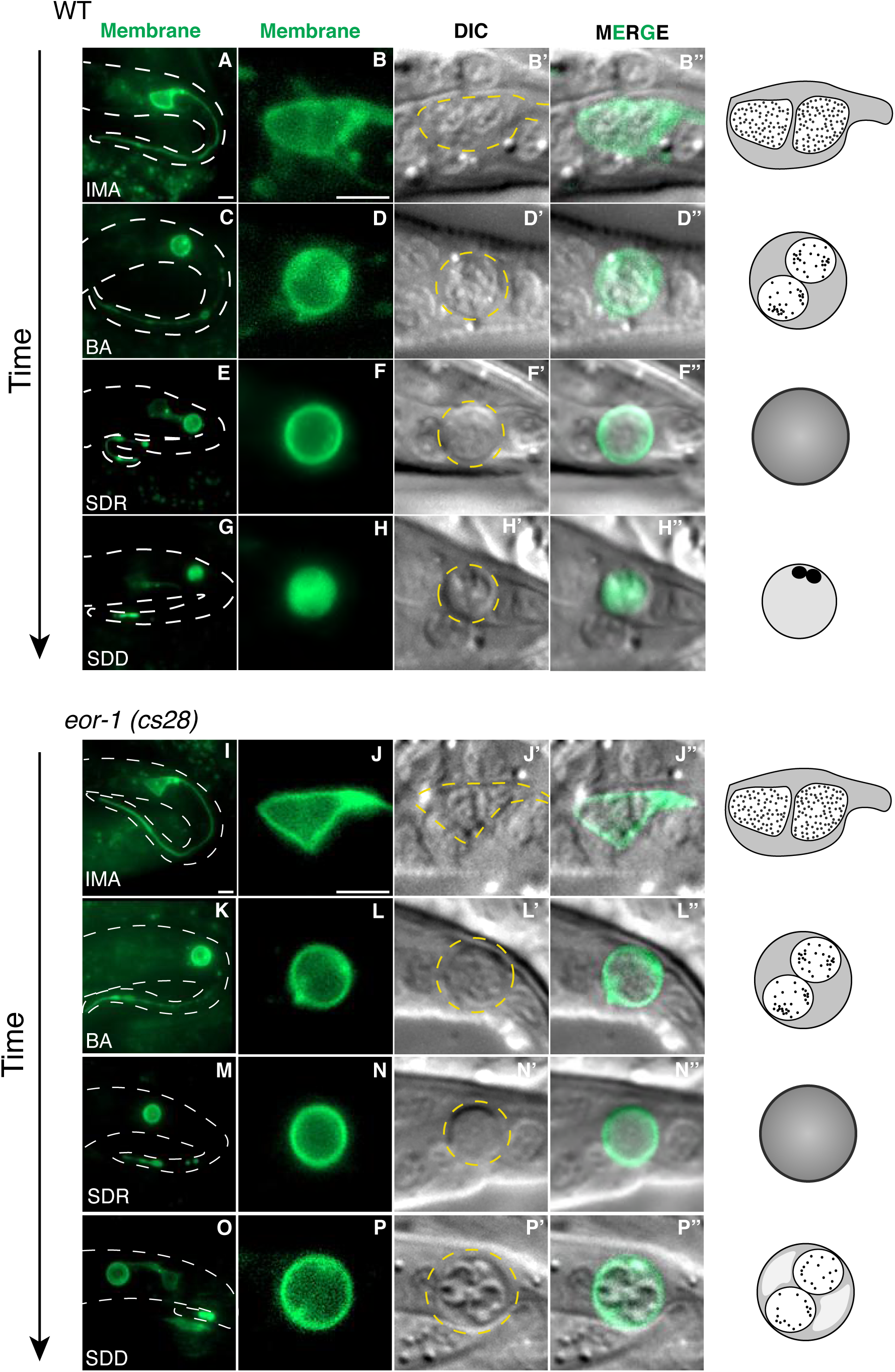
Loss of EOR-1/PLZF causes corpse arrest late in CCE progression. CCE of TSC (membrane marker) in wild-type (**A-F’’**) and *eor-1(cs28)* (**I-P’’**) mutants showing progression of soma elimination. Left-most panels show whole cell. (**A-B’’**) IMA stage in wild-type. n = 10 biologically independent animals each with similar results, (**C-D’’**) BA stage in wild-type, n = 8 biologically independent animals each with similar results, (**E-F’’**) SDR stage in wild-type, n = 10 biologically independent animals each with similar results, (**G-H’’**) SDD stage in wild-type, n = 10 biologically independent animals each with similar results. (**I-J’’**) *eor-1 (cs28)* IMA n = 8 biologically independent animals each with similar results, (**K-L’’**) *eor-1 (cs28)* BA n = 6 biologically independent animals each with similar results, (M-N’’) *eor-1 (cs28)* SDR n = 10 biologically independent animals each with similar results, (**O-P’’**) *eor-1 (cs28)* SDD n = 10 biologically independent animals each with similar results Scale bar, 5 μm.

As CCE progresses to the next stage (soma-distal process retracting stage, SDR) (**Figure 3E- F’**), the junction between the rounded soma detaches from the beading proximal process (**Figure 3E**) and the distal process starts to retract into a node. At this stage, nuclei are rounded with intranuclear contents beginning to condense into more discrete areas (**Figure 3F’-F’’**). The soma and distal process are then removed by separate phagocytes. At this stage, the soma forms a classic refractile, button-like appearance as it starts to be engulfed. After the SDR/retraction stage, the soma and process appear superficially to be degrading (Soma-distal process degrading stage, SDD) (**Figure 3G-H’’**). The soma at this point appears non-refractile and vacuolized, with its nuclei appearing pyknotic and detached **(Figure 3H’-H’’**). Apoptosis and other forms of PCD are defined by subcellular hallmarks such as specific changes in nuclear morphology and cell enlargement. We find that, during CCE, pyknotic nuclei appear also to exhibit Brownian motion within the non-refractile cytoplasm **(Figure 3H’-H’’**). The TSC membrane appears diffuse at this stage and the membrane loses its discrete ring surrounding the cell. We examined *eor-1(cs28)* TSCs in a similar manner and found that CCE progression appeared normal up until the SDD/soma-distal process degrading stage, where the nuclei fail to condense and the membrane does not lose its integrity (**Figure 3I-P’’**).

We next compared the *eor-1(ok1127)* mutant defect with that of mutants for caspase/cell killing, *ced-3(n717)* and engulfment, *ced-12(ky149),* and performed epistasis experiments. We compared the soma-persistence defects of *ced-3(n717)* (**Figure 4 A-A’’**), *ced-12(ky149)* (**Figure 4 B-B’’**) and *eor-1* mutants (**Figure 4C-C’’**). We observed that the TSC soma morphological features of *eor-1* were distinct from both *ced-3 (n717)* and *ced-12(ky149)* mutants, suggesting the cell is neither living nor engulfed. Moreover, a double mutant for *eor-1* and *ced-12(ky149)* (**Figure 4D-D’’**) phenocopied the *eor-1 (ok1127)* single mutants, rather than the refractile corpse of *ced-12(ky149)* mutants. This supports the idea that the persisting soma of *eor-1* mutants has progressed through the killing steps of CCE but has not been engulfed and is consistent with an arrest at the SDD stage. We examined whether the persisting corpse of *eor-1 (cs28)* mutants is internalized by its neighboring phagocyte. We used a cytosolic reporter labelling the TSC soma’s neighboring phagocyte (*skn-1p*::mKate2) and found that the persisting soma *eor-1* **(Figure 4E-G; Supplemental Movie S1**) mutants remains outside the phagocyt*e (*N=10/12), suggesting failure of corpse internalization.

**Figure 4.**
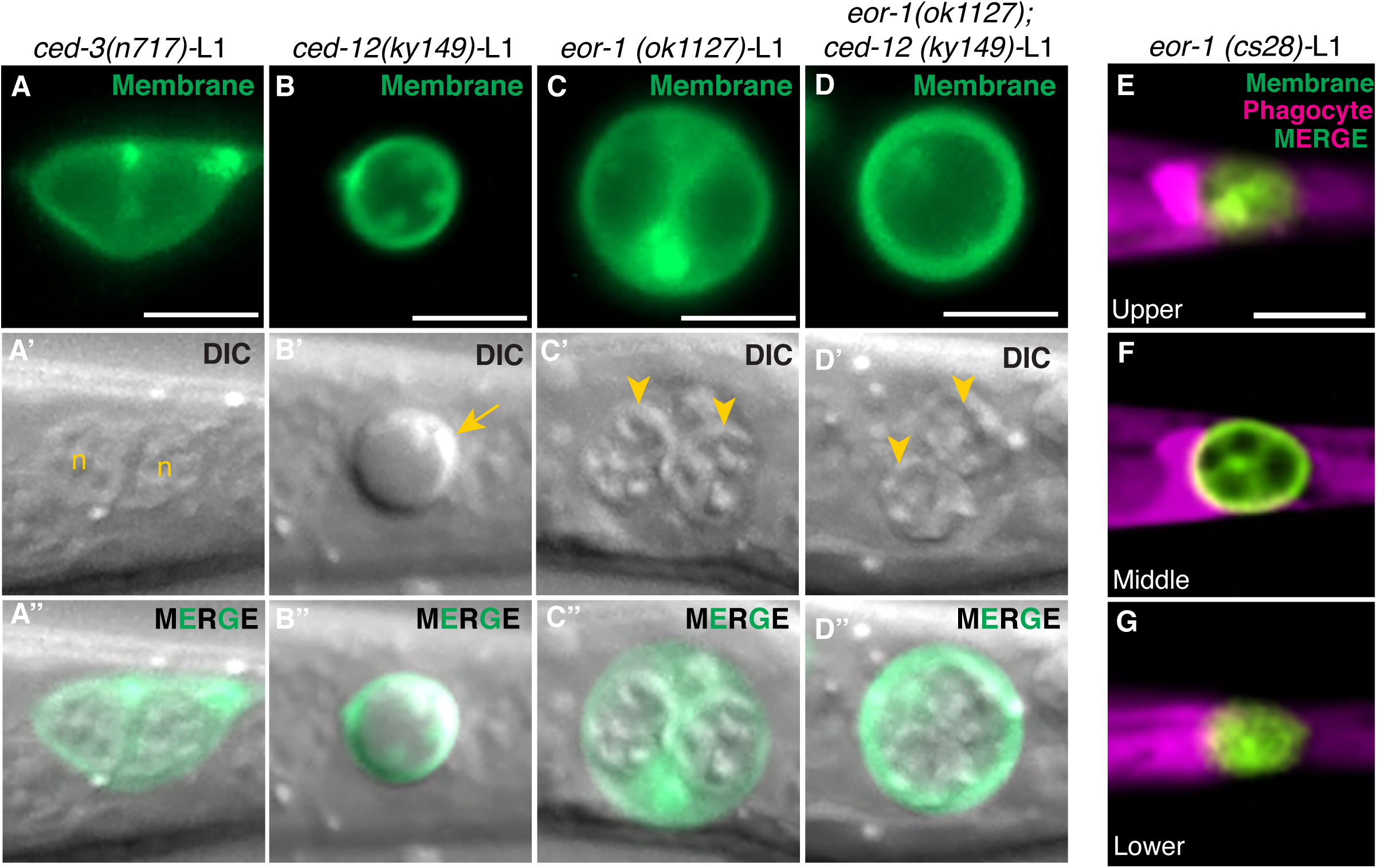
The cell corpse of *eor-1* mutants is not internalized by the neighboring phagocyte. Enlarged view of TSC soma at L1 in *ced-3(n717)* (**A-A’’**), *ced-12(ky149)* (**B-B’’**), *eor-1(ok1127)* (C-C’’), *eor-1; ced-12* (**D-D’**’), (**E-G**) Test for TSC soma corpse (green) internalization for *eor-1 (cs28)* mutant L1 *larvae* by phagocyte (magenta) at different planes of the phagocyte (upper, middle, lower). n = 10 biologically independent animals each with similar results. Scale bar, 5 μm.

### EOR-1/PLZF positively regulates *wah-1*/Aif expression in the TSC

We next sought to identify the transcriptional target for EOR-1/PLZF in the context of CCE and examined published data obtained from a previous ChiP-Seq analysis study of EOR-1/PLZF transcriptional targets (38). We noted the gene *wah-1,* which encodes a homolog of the mitochondrial flavoprotein Apoptosis Inducing Factor AIF1, a multifunctional enzyme that is essential for mitochondrial bioenergetic function (39). Interestingly, WAH-1 has been shown to function to promote apoptotic cell death in the *C. elegans* germline (40, 41). Upon activation by CED-3/Caspase and EGL-1/BH3-only, WAH-1/AIF promotes apoptosis through DNA degradation via CPS-6 endonuclease activation (41). WAH-1/AIF has also been shown to aid in phosphatidylserine (PS) exposure by promoting SCRM-1 scramblase function (40) to facilitate apoptotic corpse recognition.

We tested the hypothesis that EOR-1/PLZF promotes CCE through the transcriptional activation of *wah-1*. First, we assessed whether *wah-1* was required to promote CCE. We found that *wah-1* (*gk5392*) mutants phenocopy *eor-1* mutants (**Figure 5A, B**), and the double mutant is not additive (**Figure 5B, N=150**). We also found from cell-specific rescue experiments that *wah-1* functions in the TSC (**Figure 5C, N=150)**. We examined an endogenous reporter for *wah-1* (*wah-1*p::WAH-1::GFP) at the intact mature (IMA) embryo stage in wild-type (**Figure 5D)**, *eor- 1(cs28)* (**Figure 5E**), *mau- 2 (qm160)* (**Figure 5F)**, and *swsn-1(os22)* (**Figure 5G**) mutants and found decreased levels of *wah-1* expression in mutants (**Figure 5H**). These data support the notion that WAH-1 promotes CCE by acting within the TSC and that it is transcriptionally regulated by EOR-1, MAU-2 and SWqSN-1. This is supported by modENCODE data for *wah-1,* which identified an EOR-1 binding sequence at the *wah-1* promoter site(42). Additionally, as *wah-1* contains an HTZ-1 histone binding site, and as cohesin loaders require nucleosome-free loci to bind, our finding further implicates the importance of EOR-1 and its associated chromatin remodeling complex proteins in maintaining *wah-1* transcription during death.

**Figure 5.**
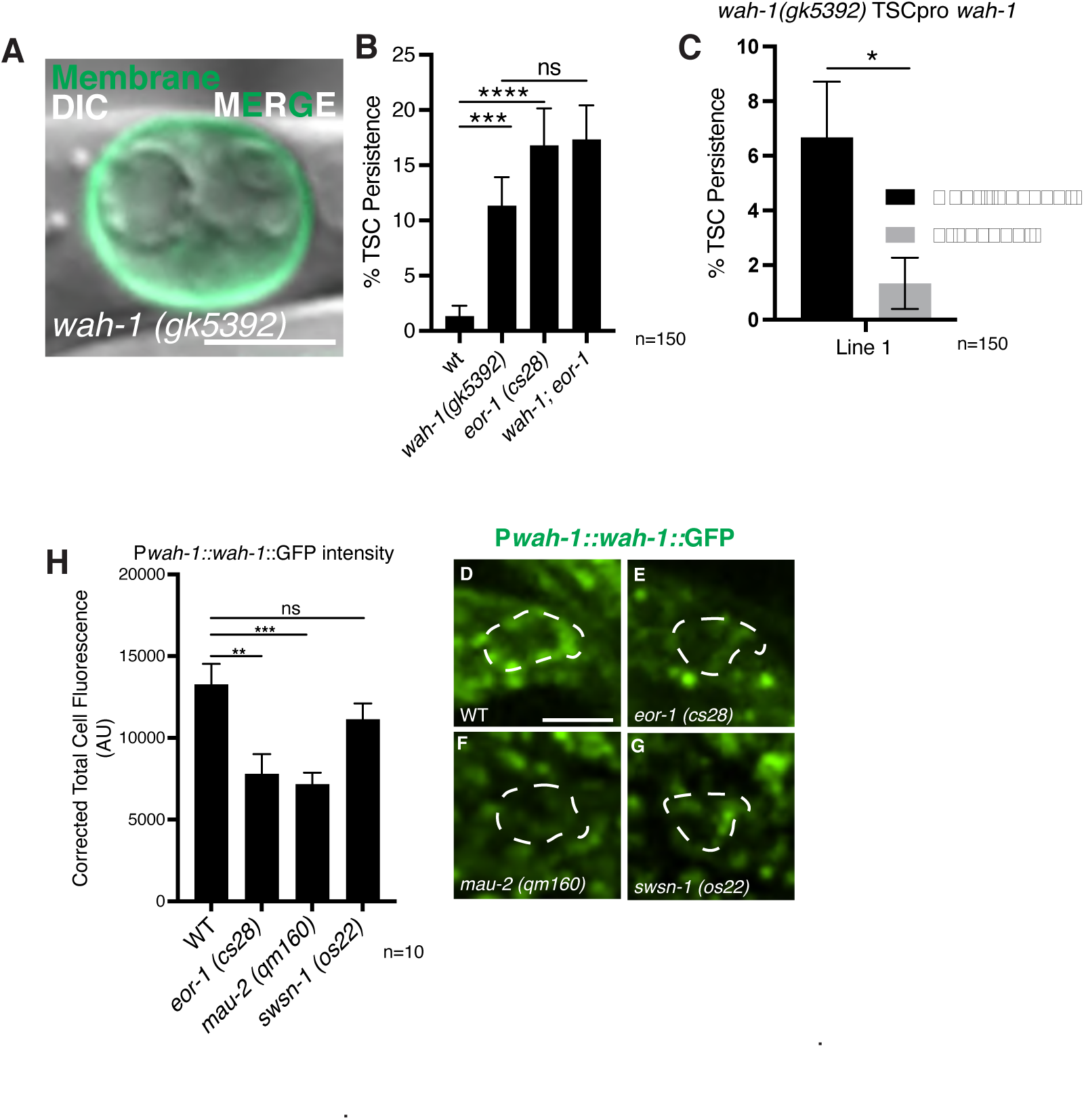
EOR-1/PLZF-1 positively regulates *wah-1*/Aif transcription to promote CCE. **(A)** *wah- 1(allele)* CCE defect of TSC at L1 n = 10 biologically independent animals each with similar results. **(B)** Quantification of *wah-1* mutant defect compared to *eor-1* and *wah-1; eor-1* double mutant. **(C)** Graph of TSC-specific rescue of *wah-1* mutant phenotype. Endogenous *wah-1* expression intensity in wild-type **(D)**, *eor-1* **(E)**, *mau-2* **(F)** and *swsn-1* **(G)** mutants. **(H)** Quantification of *wah-1* expression. n = 10 biologically independent animals each with similar results. Scale bar, 5 μm. Statistics: two-tailed unpaired student’s t-test for scoring and Mann Whittney U test for fluorescence intensity, see Supplementary Table 3 for individual P values.

### WAH-1 functions in corpse internalization

We first considered WAH-1’s possible role at the plasma membrane, as the soma rounds. Prior studies have shown that WAH-1/AIF promotes apoptotic corpse recognition and engulfment (40). We looked at CCE progression in *wah-1*(*gk5392*) (**Figure 6A-H’’**) mutants and found that the progression differed from wild-type, arresting, as in *eor-1* mutants, at the SDD stage (**Figure 6G-H’’**).

**Figure 6.**
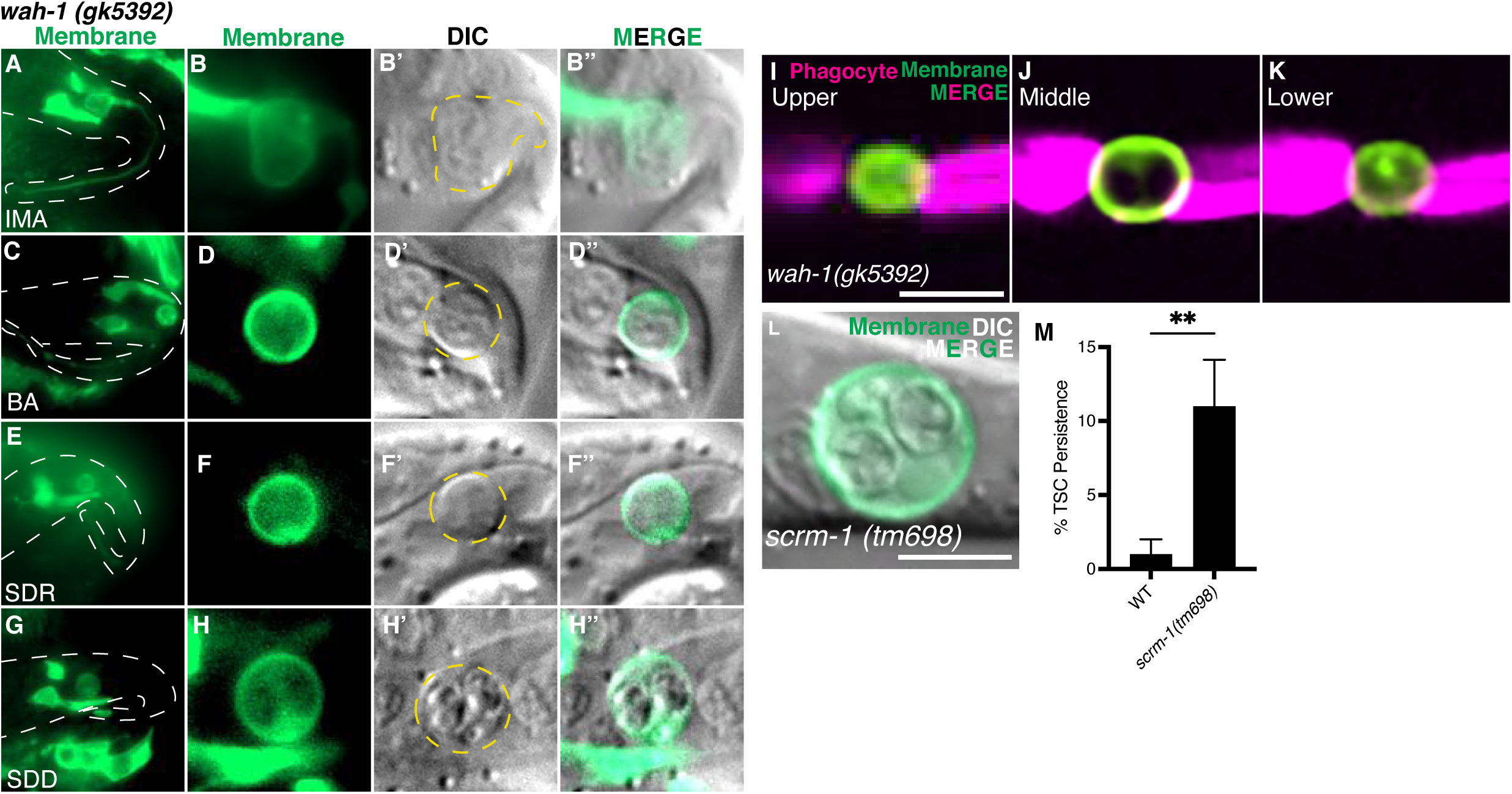
The cell corpse of *wah-1* mutants is not internalized and may function at the plasma membrane. (A-H’’) CCE progression in *wah-1(gk5392).* (**A-B’’**) IMA *wah-1 (gk5392)* n = 4 biologically independent animals each with similar results, (**C-D’’**) BA *wah-1 (gk5392)* n = 5 biologically independent animals each with similar results, (**E-F’’**) SDR *wah-1 (gk5392)* n = 4 biologically independent animals each with similar results, (**G-H’’**) SDD *wah-1 (gk5392)* n = 3 biologically independent animals each with similar results **(I-K)** TSC and soma-phagocyte dual marker strain in *wah-1* mutant L1 showing failure in TSC corpse internalization. **(L)** L1 *scrm-1* mutant showing persisting TSC soma and exaggerated nuclei. n = 10 biologically independent animals each with similar results. **(M)** Quantification of *scrm-1* mutant defect. Scale bar, 5 μm. Statistics: two-tailed unpaired student’s t-test, see Supplementary Table 3 for individual P values.

Next, we used a cytosolic reporter labelling the neighboring phagocyte (*skn-1*p::mKate2) and found that the persisting soma of *wah-1* mutants remains outside the phagocyte **(Figure 6I-K**, N=11/13), similar to *eor-1* mutants and that both *eor-1* and *wah-1* mutants the phagocyte pseudopods are not formed (**Figure 4E-G, Supplemental Movie S1; Figure 6I-K; Supplemental Movie S2**), suggesting failure in corpse recognition.

WAH-1 has been shown to promote SCRM-1 scramblase flipping of phosphatidylserine from the inner to the outer plasma membrane (40). We found that *scrm-1*(-) mutants also displayed CCE defects (**Figure 6 L, M, N=9)** akin to *eor-1* and *wah-1* mutants. These data suggest WAH-1 is important for TSC soma corpse recognition during CCE in a manner dependent on SCRM-1, as seen in its apoptotic role.

### WAH-1 also functions in nuclear dismantling after corpse internalization

WAH-1 has been shown to be involved in DNA degradation and to translocate from the mitochondria to the nucleus (41). We tested whether this is true during CCE and attempted to separate WAH-1 function in corpse internalization versus its possible nuclear role.

WAH-1 has been shown to promote DNA degradation during apoptosis by acting to enhance the activity of the CPS-6/Endonuclease G nuclease (41, 43). We found that *cps-6* mutants also phenocopy *wah-1* and *eor-1* mutants for CCE (**Figure 7A, B, N=6).** Moreover, a *wah-1; cps-6* double mutant did not show an increase in TSC defects (**Figure 7A**) suggesting these genes function in the same genetic pathway. We also found that mutants for *nuc-1*, a lysosomal acid hydrolase that promotes late DNA degradation (44, 45), also phenocopy *eor-1* and *wah-1* mutants (**Figure 7A, C)** supporting a role for WAH-1 in DNA degradation. Interestingly, it has been reported that degradation of apoptotic cell corpse DNA requires contributions of both the dying and the engulfing cell. NUC-1 is released from the apoptotic corpse into the phagolysosome, assisting in DNA degradation within the engulfing cell (45).

**Figure 7.**
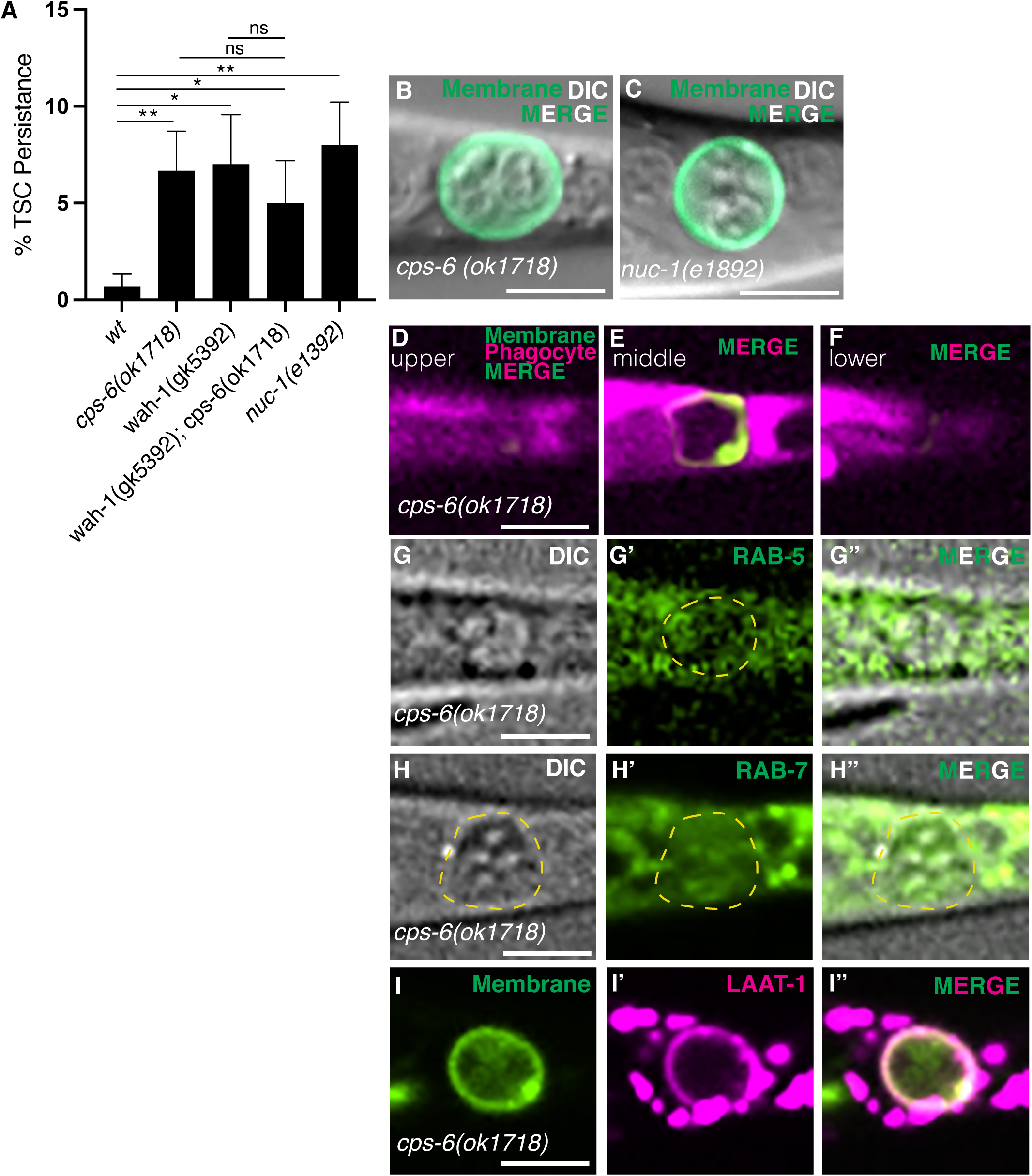
WAH-1 may promote nuclear degradation of the TSC soma corpse at phagolysosomal stage. **(A)** Quantification of *cps-6(ok1718)* and *nuc-1(e1382)* CCE defect of TSC (soma persistence) at L1. **(B)** Images of these defects. **(D)** Test for soma corpse internalization for *cps-6(ok1718)* across phagocyte planes (upper, middle lower). Localization of phagocyte GFP::RAB-5 **(G-G’’),** GFP::RAB-7 **(H- H’’)** and LAAT-1:mCherry **(I’-I’’)** relative to TSC corpse of *cps-6(ok1718)* mutant at L1 stage. n = 10 biologically independent animals each with similar results. Scale bar, 5 μm. Statistics: two-tailed unpaired student’s t-test, see Supplementary Table 3 for individual P values.

As such, we investigated whether *cps-6* mutants arrested in the phagosome maturation stage. Interestingly, unlike in *wah-1* and *eor-1* mutants, the soma of *cps-6* mutants appears to be internalized (**Figure 7D-F; Supplemental Movie S3**) (N=9/10), suggesting that CPS-6/EndoG is not involved in WAH-1’s corpse internalization function and, rather, likely plays a role in nuclear degradation downstream of internalization during phagosome maturation to resolve the internalized corpse.

To test in which stage of phagosome maturation WAH-1 and CPS-6 function together, we looked at known markers of phagosome maturation, for early (GFP::RAB-5) (**Figure 7G-G’’**), late (GFP::RAB-7) (**Figure 7H-H’’**) and phagolysosomes (LAAT-1::mCherry) (**Figure 7I-I’’**)(27) in *cps-6* mutants.We found that the phagosome bearing the internalized corpse is decorated by LAAT-1, suggesting arrest at the phagolysosomal stage; in contrast, localization of RAB-5 and RAB-7 were normal. This is consistent with a role of NUC-1 in TSC DNA degradation and supports previous hypotheses about NUC-1 being a phagolysosomal substrate. This also suggests that a defect in corpse degradation causes the dramatic nuclear and cytoplasmic phenotypes in mutants.

### Defined spatiotemporal dynamics suggest dual functions of WAH-1/AIF before and after corpse internalization

Having identified WAH-1/AIF as a regulator of CCE at both the level of corpse internalization and degradation, we examined its localization pattern across CCE stages (**Figure 8)**. First, we examined TSC mitochondrial localization against the TSC plasma membrane. Interestingly, we found mitochondria to be close to the plasma membrane through most of CCE (**Figure 8 A-D**), appearing fragmented at the SDD stage when the plasma membrane integrity is lost (**Figure 8D**). We then examined TSC promoter-driven WAH-1::mCherry against a TSC promoter-driven mitochondrial matrix marker (**Figure 8E-H’’**). When the cell is intact, WAH-1 appears to be mitochondrial (**Figure 8E-E’’**) and not in the plasma membrane. However, as the beading (BA/BD) stage begins, the soma rounds and the soma detaches from the process, both mitochondria and WAH-1 appear to be at or close to the plasma membrane (**Figure 8 F-G’’**). Afterwards, at the SDD stage, WAH-1 is not seen at the plasma membrane but can be seen in what appears to be fragmented mitochondria (**Figure 8H-H’’**); however, it is largely delocalized and cytoplasmic.

**Figure 8.**
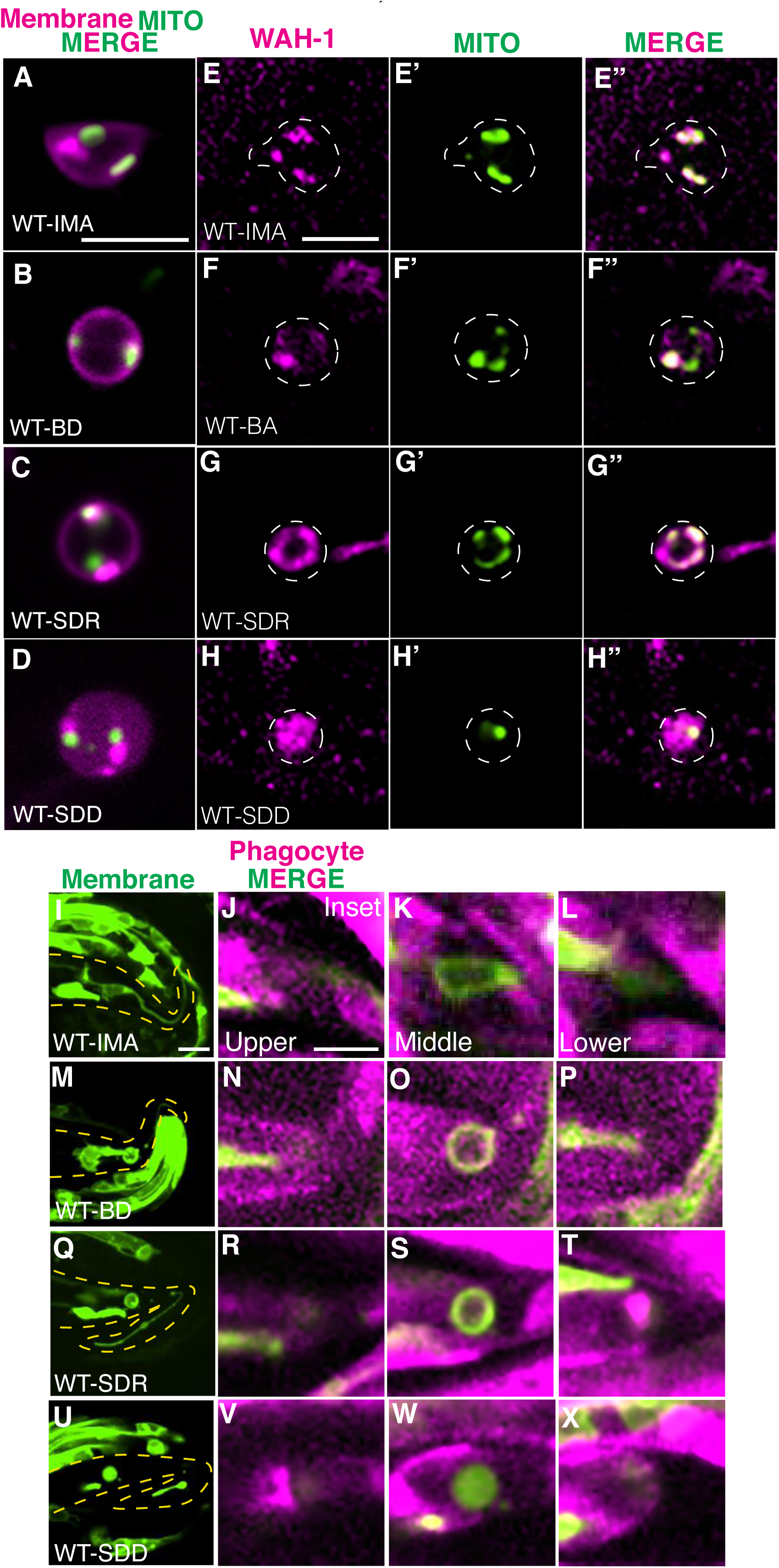
WAH-1 exhibits defined spatiotemporal dynamics during CCE progression correlated with its function. TSCp::WAH-1::mCherry and TSCp::mito::GFP localization during CCE IMA **(A**), BA (**B**), SDR (**C**) and SDD (**D**) stages in wild-type embryos. TSCp::mKate2-PH (membrane). TSCp::mito::GFP localization (different animal) during CCE IMA **(E**), BA (**F**), SDR (**G**) and SDD (**H**) stages in wild-type embryos. Relative position of TSC soma versus soma phagocyte (upper, middle and lower phagocyte planes) at different CCE stages, IMA (**I, J-L**), BD (**M, N-P**), SDR (**Q, R-T**) and SDD (**U, V-X**). n = 10 biologically independent animals each with similar results for A-D. Scale bar, 5 μm.

We imaged our TSC reporter against the soma-phagocyte reporter in wild-type embryos (**Figure 8I-X)** to examine whether the TSC soma is outside the phagocyte at the IMA stage (**Figure 8I-L)** We found that the TSC soma is internalized from what we define as the BD (beading, detached) stage when the soma rounds and appears refractile (**Figure 8M-P**) through the SDR (**Figure 8Q-T**) and SDD (**Figure 8U-X**) stages. We observe WAH-1 outside of mitochondria at the SDD stage (**Figure 8D** and **8U-X**) after being at the mitochondria and plasma membrane earlier at the SDR (**Figure 8C** and **8Q-T**) and BD (**Figure 8B** and **8M-P**) stages. Together, these data suggest that a physical association between mitochondria and the plasma membrane, possibly involving WAH-1, may be important early in phagocytosis and that exit of WAH-1 from mitochondria may be important for corpse degradation and digestion after internalization.

## Discussion

Here we describe defined subcellular events of cell corpse during phagocytosis. Our work supports a model (**Figure 9)** in which WAH-1/AIF dual functions, positively regulated by EOR-1/PLZF, ensure clearance of the TSC soma corpse during CCE. We provide the first evidence of transcriptional regulation of WAH-1. It will be interesting to explore whether the same transcriptional relationship between EOR-1 and *wah-1* exists in other forms of PCD such as LCD. We describe previously unreported morphological features of programmed cell elimination. Our study expands on prior studies linking EOR-1/PLZF to apoptotic (34) and non-apoptotic (36) PCD by providing a precise mechanistic contribution of EOR- 1/PLZF in CCE and implies that programmed cell degradation occurs in a highly regulated manner, not by just via catastrophic destruction. We also propose that regulation of DNA degradation and nuclear destruction allows the corpse to act as an active player in its own destruction, coordinating with the engulfing cell.

**Figure 9.**
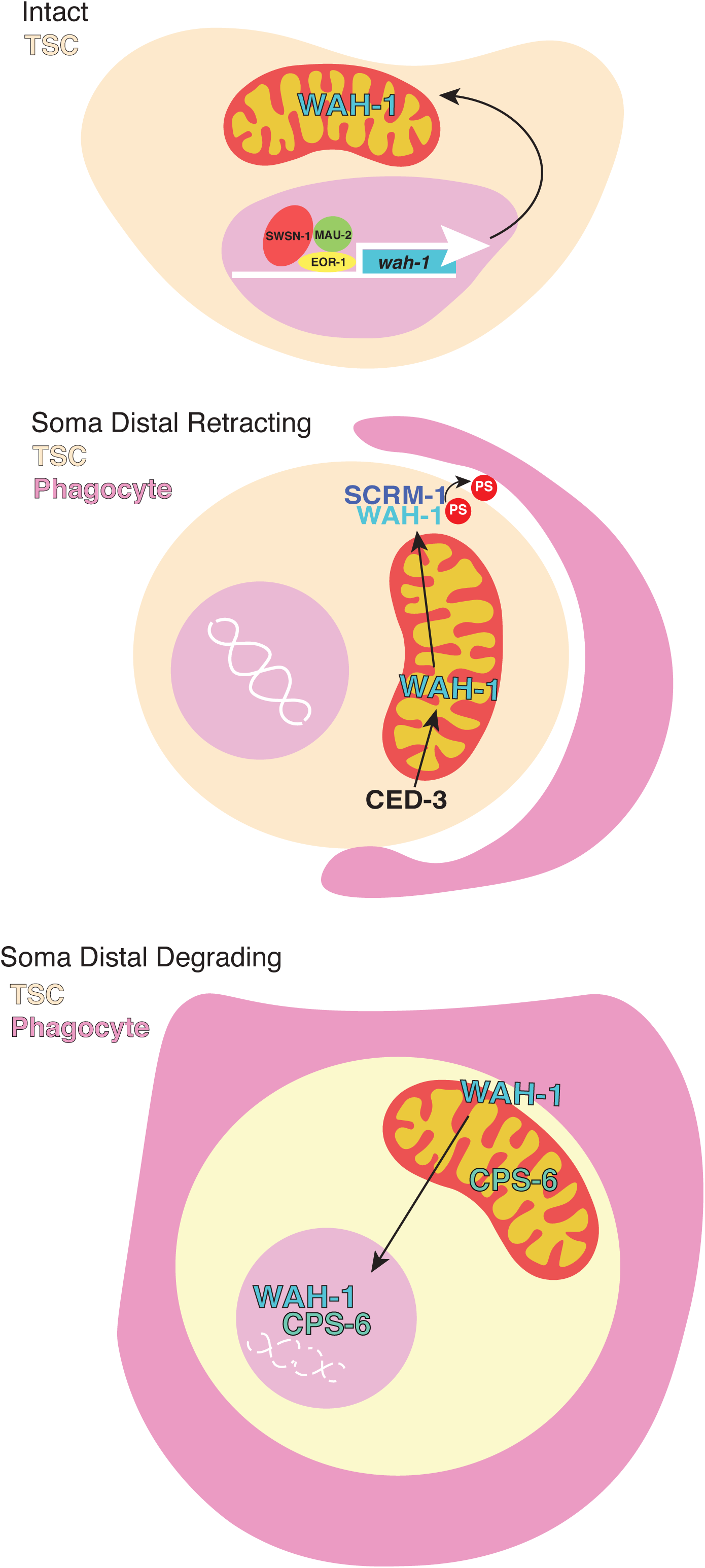
Model. EOR-1-regulated WAH-1 translocates to different TSC soma sub-compartments to sequentially promote corpse internalization and resolution.

Notably, WAH-1 exit from mitochondria is reported to be dependent on EGL-1/BH3-only (41). However, CCE is independent of EGL-1/BH3-only (30). The WAH-1 localization pattern suggests WAH- 1 can promote nuclear degradation independent of EGL-1. The lack of EGL-1 involvement may affect WAH-1’s ability to be released from mitochondria and may explain why WAH-1 is still associated to some degree with mitochondria until the SDD stage of CCE. The lack of EGL-1 may cause WAH-1’s mitochondrial release to be markedly slower, which may explain why the time course of CCE is much longer than apoptotic germline cells.

Our work also opens several exciting questions . What guides WAH-1 translocation to the plasma membrane and nucleus in a temporally regulated manner? Does phosphatidylserine triggers engulfment during CCE? What is the molecular mechanism by which WAH-1/AIF transduces the signal from the changing phagosomal environment to respond via nuclear translocation to execute its role in DNA and nuclear degradation? How does WAH-1/AIF know to translocate so specifically in the corpse at distinct phagosomal stages? We also suggest that failure to appropriately degrade DNA can lead severe exaggerations in nuclear morphology. Taken together, our study expands our understanding of the transcriptional regulation of corpse clearance and highlights roles for regulators of apoptosis in in broader contexts of programmed cell elimination.

## Materials and Methods

### C. elegans methods

*C. elegans* strains were cultured using standard methods on *E. coli* OP50 and grown at 20°C. Wild-type animals were the Bristol N2 subspecies. For most TSC experiments, one of two integrated reporters were used: *nsIs435=TSCp::myrGFP* or *nsIs685=TSCp::mKate2-PH.* In addition, *nsIs650*=*wah-1*p::WAH- 1::GFP was used. Integration of extrachromosomal arrays was performed using UV and trioxsalen (T2137, Sigma). Animals were scored at 20°C, unless otherwise mentioned.

### Imaging

Images were collected on a Nikon TI2-E Inverted microscope using a CFI60 Plan Apochromat Lambda 60x Oil Immersion Objective Lens, N.A. 1.4 (Nikon) and a Yokogawa W1 Dual Cam Spinning Disk Confocal images were acquired using NIS-Elements Advanced Research Package. For still embryo imaging, embryos were anesthetized using 0.5 M sodium azide. Larvae were paralyzed with 10mM sodium azide. Widefield imaging was performed on a Carl Zeiss Axio Imager.M2 microscope with 63X oil immersion lens. Super-resolution images were taken using a VTiSIM Super resolution Live Cell Confocal Imaging System.

### Quantifying CCE defects

TSC death defects were scored at the L1 stage. Animals were mounted on slides on 2% agarose-M9 pads, paralyzed with 10mM sodium azide, and examined on a Carl Zeiss Axio Imager.M2 microscope with 63X oil immersion lens. The persisting TSC was identified by fluorescence based on its location and morphology. At least 50 animals were scored per genotype. For images, each image is representative of 10 sampled.

### Worm strains used in this study

List gene alleles under different chromosomes linkage groups (LG I-X) LGI: *ced-12(ky149), mau-2(qm160), scrm-1(tm698), cps-6(ok1718)* LGII:

LGIII*: ced-4(n1162), wah-1 (gk5392)*

LGIV: *eor-1(ok1127), eor-1(cs28), ced-3(n717)*

LGV: *swsn-1(os22)*

LGX: *eor-2 (cs42) nuc-1 (e1392)*

### Plasmids and Transgenics

Plasmids were generated via Gibson cloning. Primer sequences and information on the construction of plasmids used in this study are provided in (**Supplemental Table 1**). The full list of transgenes is described in (**Supplemental Table 2**). The full length or fragment of the *aff-1* promoter was used to label the TSC.

### Quantification of WAH-1::GFP fluorescence intensity

Images of the intact TSC in animals harboring *wah-1* promoter driven *wah-1*::GFP in WT, *eor-1(cs28)*, *mau-2(qm160)*, and *swsn-1(os22)* were collected using confocal microscopy as stated above using the same settings across all images. Sum intensity projections of the TSC soma were generated by following TSCp::mCherry in Fiji Image J software and GFP intensity was then calculated using this software. CTCF was calculated using Microsoft Excel and graphed using Graphpad. All data points represent the average of 10 animals. Statistical significance was determined by Mann-Whitney U test for comparison between wild type and mutant animals.

### Statistics

Sample sizes and statistics were based on previous studies of CCE and the TSC(27). Independent transgenic lines were treated as independent experiments. An unpaired two-tailed *t-*test was used for all persisting TSC quantifications (GraphPad Prism). For all figures, mean ± standard error of the mean (s.e.m.) is represented. P-values are indicated in (**Supplemental Table 3**)

## Acknowledgments

We thank Dr. Jennifer Malin for comments on the manuscript. NR and PG designed the experiments and wrote the manuscript. NR, MW, KJ, RS, AE, performed the experiments. GC provided significant technical assistance. *ns957* was isolated by PG and MW in the laboratory of SS. We thank members of the Ghose lab for comments on the manuscript. Some strains were provided by the CGC, which is funded by NIH Office of Research Infrastructure Programs (P40 OD010440).

## Supplemental Tables

**Supplemental Table 1.** Plasmids used in this study.

**Supplemental Table 2.** Strains and Transgenes used in this study.

**Supplemental Table 3.** Statistics and P values.

## Supplemental Movies

Supplemental Movie S1. Test for TSC soma internalization of *eor-1 (cs28)* mutants. Green, TSC membrane, Magenta, phagocyte cytosolic marker.

Supplemental Movie S2. Test for TSC soma internalization of *wah-1(gk5392)* mutants. Green, TSC membrane, Magenta, phagocyte cytosolic marker.

Supplemental Movie S3. Test for TSC soma internalization of *cps-6(ok1718)* mutants. Green, TSC membrane, Magenta, phagocyte cytosolic marker.

