## Supplemental Table 1 for "EOR-1/PLZF-regulated WAH-1/AIF sequentially promotes early and late stages of non-apoptotic corpse removal"

**Rather et al, Plasmid Information**

| **Plasmid Number** | **Plasmid Description** | **Primers** | **Plasmid construction Details** |
| --- | --- | --- | --- |
| pPG431 | TSCp::wah-1::mCherry | Backbone FWD  GGCGCGCCgttccgaatattatgaa  Backbone REV  GGAGCATCGGGAGCCTCAGGAGC  Insert fwd- tccatactttctcatttcataatattcggaacGGCGCGCCaaaATGCTGTTGCGAGCTGTTGGAAGAC  Insert rev- ccttgctCACCATCGATGCTCCTGAGGCTCCCGATGCTCCAGCACTCTTCGCATCGTCTTCATCAC | Gibson Cloning  Backbone:  pPG401=TSCp::unc-104::linker::mCherry  Insert:  *wah-1* cDNA (synthesized) |
| pPG425 | TSCp::eor-1 cDNA | Backbone FWD  tcagaagttatttcttctacgaacgttac  Backbone REV  ataaaattttcaaatttttacatacaaatgctc  Insert FWD  gaattcattgagtaacgttcgtagaagaaataacttctgaaaaATGACATTAGTAGCTAGTTCCGAAAACAC  Insert REV  taagtatcgagcatttgtatgtaaaaatttgaaaattttatTTATTGTACATTCCACGGATTCCATGCAT | Gibson Cloning  Backbone template:  pPG270=*aff-1*p::*lnp-1* cDNA::SL2::mCherry  Insert template:  *eor-1* cDNA (synthesized) |
| pPG460 | TSCp::wah-1 cDNA | Backbone Primers as above.  Insert FWD  tccatactttctcatttcataatattcggaacGGCGCGCCaaaATGCTGTTGCGAGCTGTTGGAAGAC  Insert REV  tacctttgggtcctttggccaatcccggggatcctctagaCTAAGCACTCTTCGCATCGTCTTCATC | Gibson Cloning  pPG270=*aff-1*p::*lnp-1* cDNA::SL2::mCherry  Insert template:  *wah-1* cDNA (synthesized) |
| pPG312 | *skn-1*p::mKate2 | Backbone Primers:  FWD:  ATGGTCTCCGAGCTCATTAAcGAAAAC  REV:  GATCCTCTAGAGTCGACCTGCAGGC  Insert primers (promoter):  FWD  GAAATAAGCTTGCATGCCTGCAGGTCGACTCTAGAGGATCtgctcacagatctcaaagctgcgtgtg  REV  GCTTCATATGCATGTTTTCgTTAATGAGCTCGGAGACCATctgaaaatttggaattatttttgggaatatcg | Gibson Cloning  Backbone template  mKate2 vector  Insert template: *skn-1* genomic (fosmid) |
| pPG114 | TSCp::mito GFP_SL2_ myrmcherry | cccccggcgc gccaaaATG GCACTCCT GCAATCAC GTC  ccccccagtta actaggtgaaa gtaggatgag acaggatatc | Gibson Cloning  Backbone template:  pSM::SL2::mCherry  Inster Template:  pPG112 (AscI, EcoRV) |
| pPG112 | TSCp::mito GFP | Insert FWD  ccccccGGC GCGCCaaa ATGGCACT CCTGCAAT CACGTCTC C  Insert REV  ggggggGTC GACTTTGT ATAGTTCA TCCATGCC ATGTGTAA TC | Gibson Cloning  Backbone template:  p*glr-1*::mitoGFP (Ghose et al, 2013)  Insert template:  *aff-1* promoter 1391 bp (AscI, SalI) |
| NA | *ced-1*p::LAAT-1::mCherry | NA | Cheng et al, 2015 |
| NA | P*wah-1*::*wah-1*::GFP | NA | Wang et al, 2002 |
