## Supplemental Table 2 for "EOR-1/PLZF-regulated WAH-1/AIF sequentially promotes early and late stages of non-apoptotic corpse removal"

| **Strain name (TSC #)** | **Genotype** | **Transgene** |
| --- | --- | --- |
| OS8095 | *ced-3 (n717); nsls435* | nsIs435 = aff1p::myrGFP |
| TSC293 | *eor-1(ok1127); nsIs435* | nsIs435 = aff1p::myrGFP |
| TSC307 | *ns957; nsIs435 eor-1 fosmid (WRM0616aA08) rescue Line 1* | nsIs435 = aff1p::myrGFP |
| TSC308 | *ns957; nsIs435 eor-1 fosmid (WRM0616aA08) rescue Line 2* | nsIs435 = aff1p::myrGFP |
| TSC313 | *ns957; nsIs435 eor-1 fosmid (WRM0616aA08) rescue Line 3* | nsIs435 = aff1p::myrGFP |
| TSC363 | *unc-119(ed3) III; wgIs350(eor-1::GFP);nsIs686* | nsIs686 = pPG196 pPG196 = pSM_aff1p(small)_mKate2PHutr |
| TSC364 | *ced-12(k149);nsIs435* | nsIs435 = aff1p::myrGFP |
| TSC367 | *ced-12(k149); eor-1 (ok1127); nsIs435* | nsIs435 = aff1p::myrGFP |
| TSC376 | *mau-2(qm160); nsIs435* | nsIs435 = aff1p::myrGFP |
| TSC377 | *swsn-1(ku355); nsIs431* | nsIs435 = aff1p::myrGFP |
| TSC381 | *eor-2(cs42); nsIs435* | nsIs435 = aff1p::myrGFP |
| TSC426 | *eor-1 (cs28); nsIs435* | nsIs435 = aff1p::myrGFP |
| TSC434 | *eor-1 (cs28); nsIs528* | nsIs528 = [aff1p(4 to 2.75)::myrGFP];WT; from pPG99 |
| TSC490 | *wah-1 (gk5392); nsIs435* | nsIs435 = aff1p::myrGFP |
| TSC597 | *eor-1(cs28); wah-1(gk5392); nsIs435* | nsIs435 = aff1p::myrGFP |
| TSC602 | *eor-1(cs28); ced-4 (n1162); nsls435* | nsIs435 = aff1p::myrGFP |
| TSC614 | *cps-6 (ok1718); nsls435* | nsIs435 = aff1p::myrGFP |
| TSC615 | *nuc-1 (e1392); nsls435* | nsIs435 = aff1p::myrGFP |
| TSC627 | *pPG431; mccIs003* | pPG431=aff1p::wah-1-linker-mcherry; mccIs003=aff-1p(small)::MitoGFP |
| TSC638 | *nsEx6301 nsIs650 X; mccIs100. Line 80.10; mccIs102* | mccIs102=line 1. nsEx=Pwah-1::wah-1::GFP probably on X. mccIs100=mccEx207=aff-1p(small)::TSC myrmCherry |
| TSC655 | *eor-1 (cs28); nsEx6301 nsIs650 X; mccIs100. Line 80.10; mccIs102* | mccIs102=line 1. nsEx=Pwah-1::wah-1::GFP probably on X. mccIs100=mccEx207=aff-1p(small)::TSC myrmCherry |
| TSC658 | *eor-1(cs28); ppg425; nsls435; mccEx265* | mccEx265= pPG146 =aff1p::eor-1 |
| TSC659 | *eor-1(cs28); ppg425; nsls435; mccEx266* | mccEx265= pPG146 =aff1p::eor-1 |
| TSC660 | *eor-1(cs28); ppg425 5.0ng; nsls435; mccEx267* | mccEx265= pPG146 =aff1p::eor-1 |
| TSC670 | *eor-1 (cs28); nsIs435;mccEx096* | mccEx096=pPG312: 20ng/uL, myo-2GFP: 5ng/uL, pBSK: 80ng/uL; nsIs435=aff-1p(small)myrGFP. skn-1 expression (pPG312 injected into OS8095. pPG312: 20ng/uL, myo-2GFP: 5ng/uL, pBSK: 80ng/uL) |
| TSC672 | *mau-2(qm160); nsEx6301 nsIs650 X; mccIs100. mccIs102* | mccIs102=line 1. nsEx=Pwah-1::wah-1::GFP probably on X. mccIs100=mccEx207=aff-1p(small)::TSC myrmCherry |
| TSC673 | *swsn-1(os22); nsEx6301 nsIs650 X; mccIs100.mccIs102* | mccIs102=line 1. nsEx=Pwah-1::wah-1::GFP probably on X. mccIs100=mccEx207=aff-1p(small)::TSC myrmCherry |
| TSC676 | *scrm-1(tm698); nsls435* | nsIs435 = aff1p::myrGFP |
| TSC678 | *cps-6 (ok1718); wah-1 (gk5392); nsls435* | nsIs435 = aff1p::myrGFP |
| TSC680 | *wah-1(gk5392; pPG460 (wah-1 rescue) 1.0ng line 2; nsls435; mccEx275* | pPG460=aff1p::wah-1 mccEx275 nsIs435 = aff1p::myrGFP |
| TSC694 | *cps-6 (ok1718); nsIs435;mccEx096* | mccEx096=pPG312=skn-1 expression mKate2 nsIs435=aff-1p(small)myrGFP. |
| TSC695 | *cps-6 (ok1718); nsIs435;nsEx5971* | nsEx5971 = ced-1p::LAAT-1::mCherry + coel-GFP nsIs435 = aff1p::myrGFP +coel-red |
| TSC696 | *cps-6 (ok1718);rab-7(utx12[mNG::rab-7])* | Superficially wild-type. N-terminal tag of RAB-7 via CRISPR/Cas9 knock-in of mNeonGreen at rab-7 locus. Insertion verified by PCR. Left flank: 5' gcacaacaaaaaggcttccagtgaacaaaa 3'; Right flank: 5' ATGTCGGGAACCAGAAAGAAGGCGCTGCTC 3'. sgRNA: 5' cttccagtgaacaaaaATGT 3'. CRISPR/Cas9 homologous recombination, not outcrossed. |
| TSC698 | *wah-1(gk5392); nsIs435;mccEx280* | mccEx280=pPG312=skn-1 expression mKate2 nsIs435=aff-1p(small)myrGFP. |
| TSC703 | *nsIs435;mccEx280* | mccEx096=pPG312=skn-1 expression mKate2 nsIs435=aff-1p(small)myrGFP. |
| TSC705 | *rab-5(udn14) I; udnSi38; cps-6 (ok1718)* | udnSi38 [rab5p::rab-5] II. rab-5 [D135H]. Homozygous lethal rab-5 [D135H] mutation rescued by a single copy of wild-type rab-5 integrated into chromosome II at ttTi5605 site (II: 0.77). |
