## Supplemental Table 3 for "EOR-1/PLZF-regulated WAH-1/AIF sequentially promotes early and late stages of non-apoptotic corpse removal"

| **Figure Panel** | **Figure 1** | **P value** |
| --- | --- | --- |
| 1F | wt 20°C vs wt 25°C | 0.3198 |
| 1F | wt 20°C vs n2957 20°C | <0.0001 |
| 1F | wt 25°C vs n2957 25°C | <0.0001 |
| 1I | *ns957* fosmid rescue line 1 non-transgenic vs transgenic | <0.0001 |
| 1I | *ns957* fosmid rescue line 2 non-transgenic vs transgenic | <0.0001 |
| 1I | *ns957* fosmid rescue line 3 non-transgenic vs transgenic | <0.0001 |
| 1J | wt 25°C vs eor-1 (ok1127) | <0.0001 |
| 1J | wt 25°C vs eor-1 (cs28) | <0.0001 |
| 1L | *eor-1(cs28)* pTSC::EOR-1 rescue Line 1 non-transgenic vs transgenic | 0.0012 |
| 1L | *eor-1(cs28)* pTSC::EOR-1 rescue Line 2 non-transgenic vs transgenic | 0.0294 |
| 1L | *eor-1(cs28)* pTSCTSC::EOR-1 rescue Line 3 non-transgenic vs transgenic | 0.0008 |
|  | **Figure 2** |  |
| 2A | wt 25°C vs *eor-1 (ok1127)* | <0.0001 |
| 2A | wt 25°C vs *eor-1 (cs28)* | <0.0001 |
| 2A | wt 25°C vs *eor-2 (cs42)* | 0.0056 |
| 2A | wt 25°C vs *mau-2 (qm160)* | <0.0001 |
| 2A | wt 25°C vs *swsn-1 (ku355)* | 0.0017 |
|  | **Figure 5** |  |
| 5B | wt 25°C vs *wah-1 (gk5392)* | 0.0003 |
| 5B | wt 25°C vs *eor-1 (cs28)* | <0.0001 |
| 5B | *eor-1 (cs28)*; *wah-1 (gk5392)* vs *eor-1 (cs28)* | 0.9073 |
| 5B | *eor-1 (cs28)*; *wah-1 (gk5392)* vs *wah-1 (gk5392)* | 0.139 |
| 5C | *wah-1(gk5392)* pTSC::WAH-1 rescue Line 1 non-transgenic vs transgenic | 0.0184 |
| 5H | Pwah-1::GFP wt vs *eor-1 (cs28)* | 0.0021 |
| 5H | Pwah-1::GFP wt vs *mau-2 (qm160)* | 0.0003 |
| 5H | Pwah-1::GFP wt vs *swsn-1 (os22)* | 0.2475 |
|  | **Figure 6** |  |
| 6M | wt 25°C vs *scrm-1 (tm698)* | 0.0028 |
|  | **Figure 7** |  |
| 7A | wt 25°C vs *cps-6 (ok1718)* | 0.0056 |
| 7A | wt 25°C vs *nuc-1 (e1392)* | 0.0017 |
| 7A | *cps-6 (ok1718)*; *wah-1 (gk5392)* vs *cps-6 (ok1718)* | 0.5885 |
| 7A | *cps-6 (ok1718)*; *wah-1 (gk5392)* vs *wah-1 (gk5392)* | 0.5538 |
